# SPECTRE: A versatile data framework for integrating spectral, environmental, and ecophysiological trait data

**DOI:** 10.64898/2026.11.22.760282

**Authors:** Vincent P. Riedel, Sylvain Colomer, Emilie Isasa, Justine Gagne, Nicholas Provart, Ian Major, Ingo Ensminger

## Abstract

With the increasing availability of large datasets and the growing importance of artificial intelligence (AI), robust and sustainable data management systems have become essential for plant and ecosystem research. In particular, the integration of heterogeneous spectral data with plant functional traits and environmental measurements from multiple sensors poses a major challenge, requiring significant time investment and long-term planning. Here, we present SPECTRE (**Spec**tral, **T**rait, and **R**emote-sensing **E**cophysiology data framework), a versatile data framework based on simple, open data formats that addresses the need of many research groups for a standardized organization of heterogeneous datasets, facilitating their integration and joint analysis without relying on proprietary software or external service providers. SPECTRE integrates a wide range of datasets linking spectral measurements to plant traits, including LiDAR, hyperspectral imagery, chlorophyll fluorescence, geospatial data, soil moisture, climate variables, and ecophysiological traits, organized into three main categories: (i) unmanned aerial vehicle (UAV)-acquired data, (ii) ground-based trait measurements, and (iii) environmental data. All datasets are stored using a standardized folder structure and converted to comma-separated values (CSV) format. A unified primary key enables consistent linking of spectral signatures to functional traits across datasets by species and site, facilitating trait-based analyses, data framework extension, and long-term usability. The widespread use of lookup tables reduces data redundancy and facilitates the integration of data with different naming conventions, while the use of Markdown for metadata and documentation ensures easy access to information without relying on complex or proprietary formats, simplifying documentation creation and maintenance. Overall, SPECTRE offers a simple, transparent, and scalable solution that can be readily adopted by plant ecophysiological laboratories, facilitating future AI-driven trait prediction analyses.

## 1. Introduction

Forest ecosystems cover 31% of the earth’s terrestrial area and account for 3.53 ± 0.41 Pg C yr⁻¹ sequestration (FAO, 2020; Pan et al., 2024). However, the latest observations indicate a high vulnerability of terrestrial ecosystems functioning as a carbon sink under climate change (Ke et al., 2024). Mitigating these adverse effects and advancing proactive strategies, such as climate-smart forestry, requires a paradigm shift in environmental monitoring (Gessler et al., 2026; Groover et al., 2025). The development of high-performance, affordable Unmanned Aerial Vehicles (UAVs), as well as the miniaturization of powerful sensors, has profoundly transformed vegetation monitoring (Syed Hanapi & Johari, 2019). In addition, the advent of artificial intelligence (AI) and deep learning allows for large datasets to be processed much more quickly, which enables the discovery of new solutions. However, standardization of data and metadata is necessary for effective data analysis (Hart et al., 2016; Wilkinson et al., 2016), particularly as learning-based approaches increasingly rely on large volumes of data, whether to train or to adapt models to specific applications. Here we introduce the SPECTRE (**Spec**tral, **T**rait, and **R**emote-sensing **E**cophysiology) data storage framework, which aims to be a low-cost, easy-to-implement, standardized, and versatile solution, created for every forestry remote sensing laboratory and beyond.

Over the past decade, forest monitoring capabilities have been transformed by rapid developments in remote sensing (RS) technologies. In particular, the deployment of high-performance, cost-effective Unmanned Aerial Vehicles (UAVs), alongside spaceborne satellite constellations, has enabled high-frequency data collection across vast spatial scales with significantly reduced labor intensity (Syed Hanapi & Johari, 2019). Rapid technological advances and growing research networks have dramatically increased the scale, diversity, and complexity of scientific data, ranging from handwritten lab notebooks to petabyte-scale databases (Hampton et al., 2013; Reid et al., 2014). Multi-sensor remote sensing (e.g. capturing LiDAR structural metrics, hyperspectral reflectance, and solar-induced chlorophyll fluorescence) provides non-destructive proxies for canopy morphology, biochemical composition, and physiological performance (D’Odorico et al., 2020; Harikumar et al., 2022; Yang et al., 2026). Furthermore, the exceptional precision of this technology enables accurate differentiation between plant diseases and drought stress, making it unavoidable for the future (Ensminger, 2020; Fallon et al., 2020). However, remote sensing imagery alone cannot fully resolve underlying physiological mechanisms. To improve model accuracy and interpretation, remote sensing observations must be regularly calibrated and validated against ground-based, species-specific ecophysiological trait measurements (e.g., leaf water potential, photosynthetic efficiency) (Cavender-Bares et al., 2020; Porcar-Castell et al., 2021).

Concurrently, the advent of artificial intelligence (AI) and deep learning has expanded our capacity to process massive, high-dimensional ecological datasets rapidly, offering new ways for precision decision-making in forest management (Eli-Chukwu, 2019; Huth et al., 2025; Colomer et al., 2026). Machine learning (ML) models have the power to predict phenotypic traits such as yield, biotic, and abiotic stress tolerance and that because of the way they capture the non-linear relationships between the genetic variation and environmental features extracted from the dataset (Danilevicz et al., 2022). For tree species mapping and for management purposes especially the individual-tree crown segmentation (ITCS), which requires complex algorithms, is a promising tool as the time-consuming task of tree delineation could be automated and the individual tree contribution or behavior can be detected (Colomer et al., 2026). Similarly, machine learning models can exploit spectral observations to predict ecophysiological traits, providing valuable insights into vegetation functioning. And ultimately, when enough data and computational power are available, large-scale models such as foundation models can be trained to learn generalizable representations, enabling them to be adapted to a wide range of tasks and opening new opportunities for more advanced decision-making in forest management (Müller et al., 2019; Hong et al., 2023; Siméoni et al., 2025).

Fully realizing the potential of multi-sensor integration and AI for rapid forest monitoring fundamentally depends on access to sufficient, high-quality training data. Furthermore, there is a necessity of diverse training datasets to minimize the sampling biases and enhance model robustness (Arndt et al., 2024; Cherif et al., 2025). However, acquiring annotated datasets for deep learning models presents a persistent bottleneck in forestry applications (Padarian et al., 2019). Unlike commercial or social media domains that harvest massive volumes of passively labeled data (Deng et al., 2009), forestry research relies heavily on resource-intensive field inventories and remote sensing observations that still demand expert manual interpretation and curation (Kangas et al., 2018; Reichstein et al., 2019). This scarcity of training data is further compounded by “data entropy”, which is the gradual loss of data accessibility over time caused by neglected long-term storage infrastructure, missing metadata, or insufficient documentation. (Michener et al., 1997; Pepe et al., 2014; Vines et al., 2014). To prevent data decay and ensure that multi-sensor datasets remain accessible for ongoing AI-driven modeling, researchers must implement proactive data management plans across the entire data lifecycle (Michener, 2015).

For now, most of the databases in the field of plant ecology/ecophysiology and remote sensing of the environment focus on field data, but rarely combine data from the UAV, ground data, and environmental data. Plant functional trait and flora repositories, such as the Ecological Flora of the British Isles (Fitter & Peat, 1994), the Seed Information Database (Liu et al., 2019), BIOPOP (Poschlod et al., 2003), GLOPNET (Wright et al., 2004), LEDA (Kleyer et al., 2008), BROT (Paula et al., 2009), USDA PLANTS data (Green, 2009), and BRIDGE (Baraloto et al., 2010), are structured primarily as relational or flat tabular databases indexed by unique taxonomic identifiers. In these architectures, species-level primary keys link standardized matrices of morphological, physiological, and life-history traits directly to categorical ecological attributes, geographic sampling locations, and measurement protocol metadata. However, these databases are either specialized for geographic regions (LEDA, BIOPOP, BROT, USDA Plants, Ecological Flora of the British Islands, BRIDGE) or specific traits (GLOPNET, SID). There are also global databases available like ‘*TRY – A plant trait database’* database (Kattge et al., 2011) or a recently developed database for whole plant hydraulic conductance KPLANT (Zhao et al., 2026). Remote sensing and integrated ecological observatory databases, such as the Forest Observation System (Schepaschenko et al., 2019), the National Ecological Observatory Network (NEON; Kampe et al., 2010; Thorpe et al., 2016), the Integrated Carbon Observation System (ICOS; Heiskanen et al., 2022), the Australian SuperSite Network (TERN; Karan et al., 2016), the Chinese Ecosystem Research Network (CERN; Fu et al., 2010), FLUXNET (Pastorello et al., 2020), OpenForest (Ouaknine et al., 2025), and NASA’s AfriSAR (Fatoyinbo et al., 2021), are structured primarily as relational models, spatial catalogs, or hierarchical array formats (HDF5, NetCDF) indexed by unique spatial coordinates. In these architectures, plot- or site-level primary keys link ground inventory measurements, flux time series, and field attributes directly to co-registered airborne or satellite imagery, LiDAR point clouds, and campaign metadata. However, despite these advancements, there is currently a lack of unified frameworks that systematically integrate remote sensing observations with fine-scale plant ecophysiology (such as leaf water potential or photosynthetic quenching parameters) and site-specific microenvironmental data within a single standardized structure.

To resolve these challenges, we introduce SPECTRE (Spectral, Trait, and Remote-sensing Ecophysiology), a lightweight, versatile, and open-access data framework engineered specifically for forestry remote sensing and ecophysiology research. SPECTRE departs from complex database engines by adopting a transparent, file-and-folder architecture organized around a standardized hierarchical directory structure. It relies exclusively on open-source, non-proprietary formats (Markdown (MD), Comma Separated Value (CSV)). Furthermore, SPECTRE enables a direct link of study sites with ITCS masks so that future data can be directly used and integrated crosslinked between ground and UAV and all that at a fraction of the time with low cost. SPECTRE facilitates long-term data preservation and usability by limiting the need for continuous software maintenance and dedicated server infrastructure. The framework categorizes data into three core domains: (1) UAV-acquired spectral imagery and spatial layers, (2) ground-based ecophysiological and morphological trait measurements, and (3) environmental data. Through a unified primary key tagging system, SPECTRE establishes explicit relational links across species identities, individual tree observations, environmental parameters, and spectral signatures across scales.

The objective of this paper is twofold:

1. To describe demonstrate how this structured framework simplifies multi-scale spectral-trait integration, facilitates cross-institutional collaboration, and prepares raw ecophysiological data for ingestion into specialized AI models to support climate-smart forestry and sustainable ecosystem management.
2. To present SPECTRE as a low-cost, adaptable data management protocol that standardizes folder organization, file naming conventions, and metadata structures for individual ecological and remote sensing laboratories.

## 2. Concept

The SPECTRE data framework is specialized in forest spectral data (remotely sensed or directly measured), environmental data and ecophysiological traits. SPECTRE is built around a standardized hierarchical folder structure, supported by open data formats and lightweight metadata documentation. Its purpose is to make data analysis efficient and thereby improving data management in individual labs as well as making collaboration more effective. Therefore, this data framework follows the principles shown in Figure 1.

**Figure 1:**
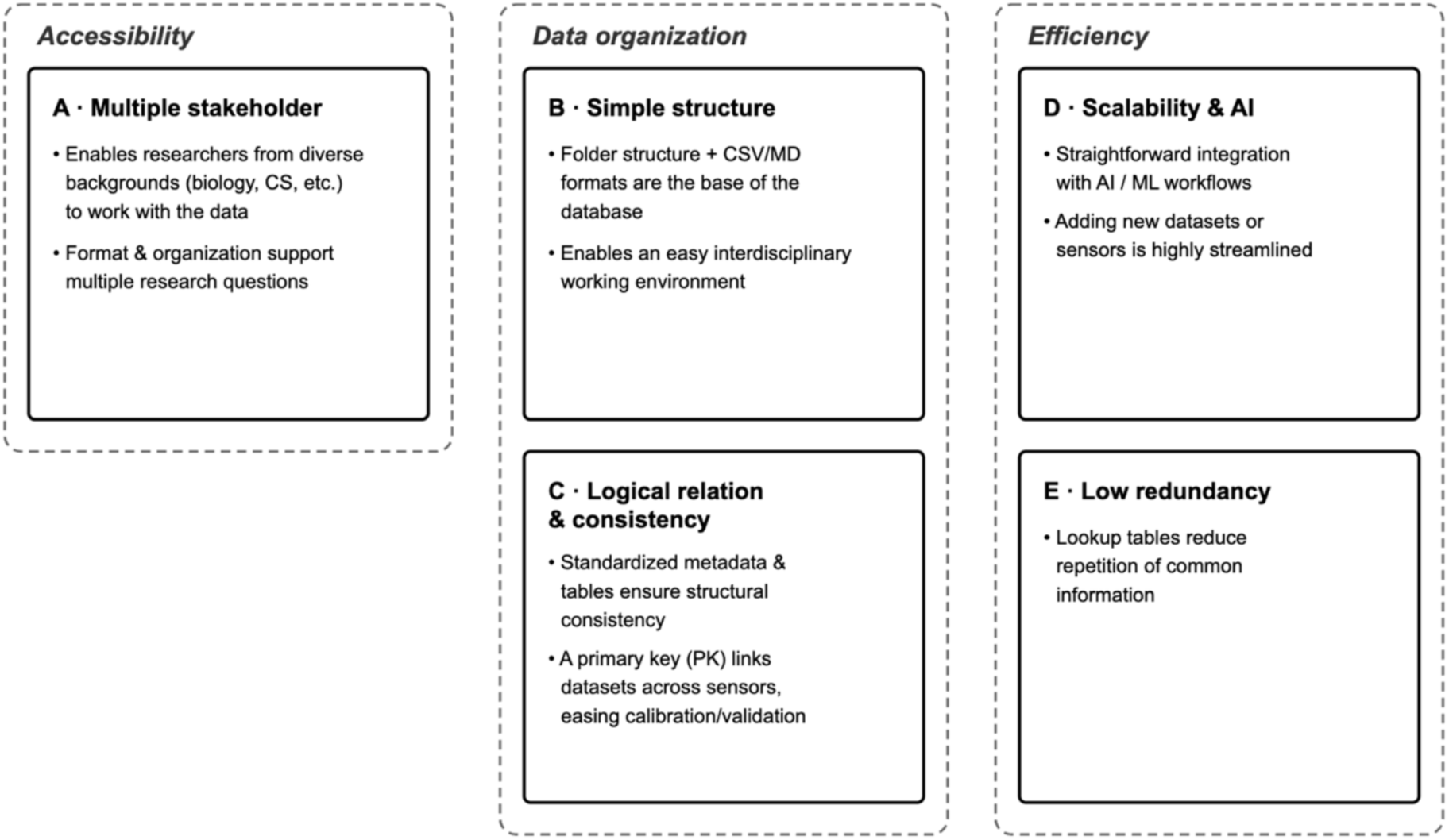
The principles of the SPECTRE data framework. The principles are grouped by overall themes by the dashed lines

Overall, these principles provide a straightforward approach to introduce data organization and long-term storage to a wide range any research group or collaboration.

## 3. Implementation

The following describes the data model, data formats, the technology and the interface of the SPECTRE data framework. The development is based on the previously mentioned principles. A basic layout is shown in Figure 2, demonstrating individual steps and authorities from data acquisition to the user interface. Abbreviations are explained in Table S1.

**Figure 2:**
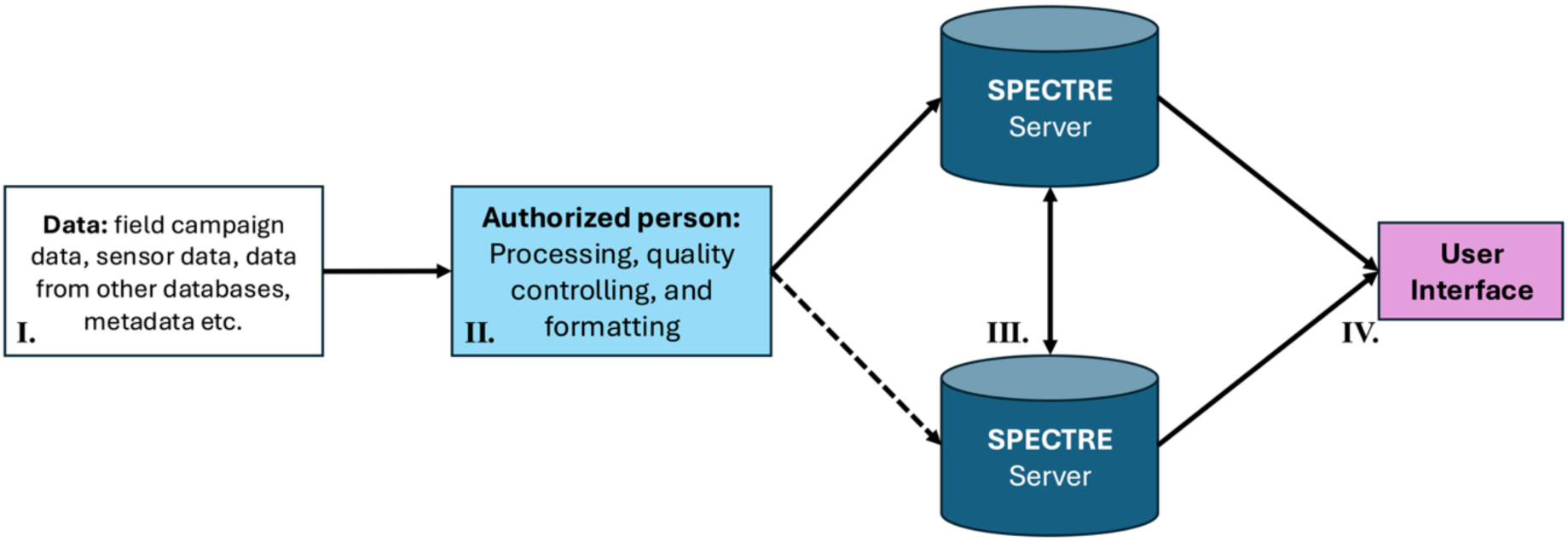
Basic layout of the SPECTRE data framework. **I.** Data retrieval from different sources. **II.** The control and processing by an authorized person. **III.** Two physically and geographically separated mirroring storage server. **IV.** A simple web-based user interface allows the search and download of the datasets.

### 3.1 Data model

The data model determines the structure of data and within the data framework SPECTRE organizes the data in a combination of a hierarchical folder structure with a relational model structure (Figure 3 and Figure 4). This combination allows for a simple data organization, while keeping it versatile and scalable.

**Figure 3:**
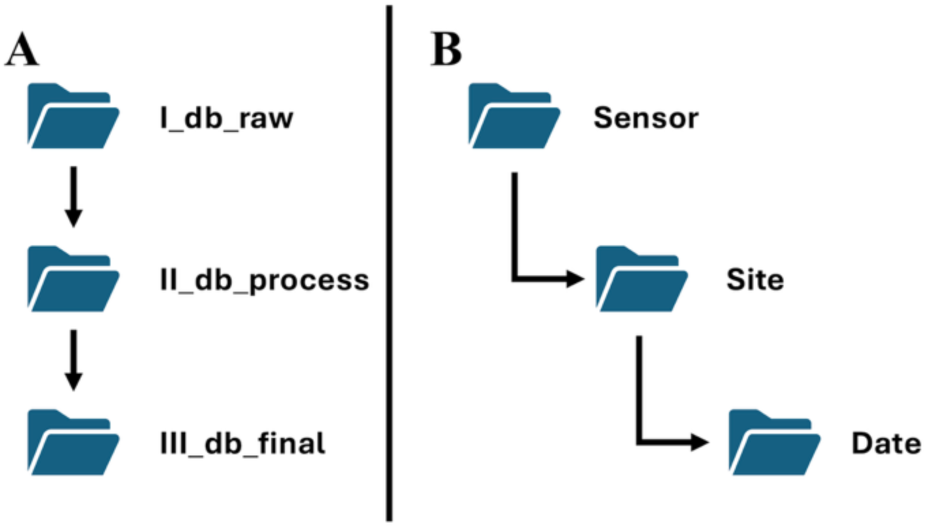
Hierarchical data model of the SPECTRE data framework. **A.** Copy of three identical folder structures for raw data (I_db_raw), processed (II_db_process), and final datasets (III_db_final). **B.** Hierarchical folder structure within each of the three identical folder structures.

**Figure 4:**
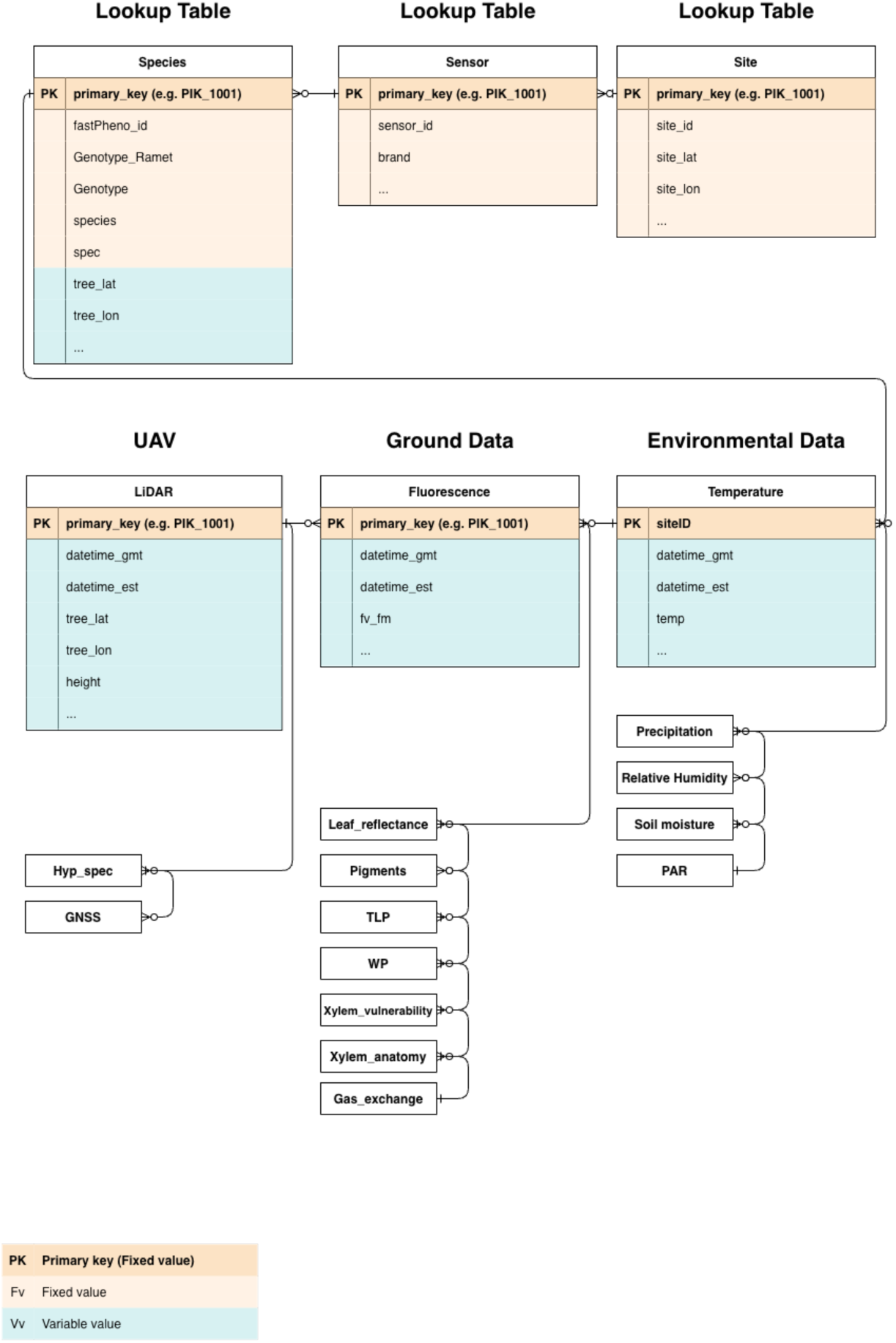
Entity-Relationship-Diagram (ERD) of the SPECTRE data framework, which describes the relationship between the datasets. Datasets are grouped into three major groups: UAV imaged, Ground measured, and Environmental Data. Furthermore, there are lookup tables for reducing the redundancy. The orange marked fields are columns within a dataset which contains fixed information, like a certain tree or site. Also, the primary key is a fixed value, and its purpose is the linking of all datasets throughout the data framework. The blue fields are variable columns in a which mostly the sensor measured values are represented, or the time for example.

#### 3.1.1 Hierarchical folder structure

The SPECTRE data framework is specialized for forest remote sensing and ecophsysiological data. To ensure that data are organized and stored as raw-data, processed-data, and final-data SPECTRE is built upon three main folders with an identical hierarchical folder structure (Figure 3, B.). Therefore, the three folders are called ‘*I_db_raw*’, ‘*II_db_process*’, and ‘*III_db_final*’ (Figure 3, A.). Each folder has a distinct purpose and access permissions: *I_db_raw* is private with restricted access to protect the original data, *II_db_process* serves as intermediate storage for data processing, and *III_db_final* is publicly accessible online and serves as the reference repository for finalized datasets. While the basic folder structure is identical within each of the three folders, data formats of the datasets can vary due to sensor-specific data format retrieving, processing, and transforming of the data (see Section 3.2).

#### 3.1.2 Entity relationship model

The hierarchical folder structure of the SPECTRE framework includes a relation between the different datasets through the establishment of a primary key (PK), similar to that used in relational databases (see Figure 4). Therefore, the datasets themselves are grouped into three major sub-groups UAV, Ground measured, and Environmental.

Before explaining the separation of the datasets into three groups it is important to understand the link between the datasets. SPECTRE data framework use PKs, which combines a site information with a tree information (mostly in the form of increasing integers). Due to this combination each tree gets a unique name. By giving every tree a unique name, measurements from various sensors measured at various distances to the tree can be linked, facilitating their joint analysis.

The UAV/Remotely sensed group includes datasets which are received from sensors flying above the ground (e.g. UAV, plane, satellite). This group is from a storage perspective a group with the largest amount of data.

The Ground measured group includes all data measured by hand in the field or laboratory with all kinds of tree spectral/ecophysiological sensors (e.g. FluorPen, UniSpec, HPLC, Scholander Pressure Chamber). Separating this group from UAV is done not only because there is a variety of traits which can’t be measured with a UAV, but also because these measurements can be used for calibration and validation of the UAV imaged spectral data.

#### 3.1.3 Reduced redundancy through lookup tables

As a further structural element lookup tables (LKT) have been created. These have the main function of storing repetitive information within one single table. Repetitive information means information, which is independent from a main group or sensor, and which would create unnecessary storage usage. For example, genotype information for a single tree is a unique information, that doesn’t change. Instead of integrating this information into every column of every trait it is less redundant to keep this information in a separate LKT. For specific research questions this genotype information can be then linked to the required traits by the primary key (PK) (see Section 3.1).

#### 3.1.4 Metadata

Metadata are an important documentation tool giving each dataset a long-lasting documentation and, for all data users, the possibility of a correct interpretation and analysis of the data. For the SPECTRE data framework each dataset has a corresponding metadata in the Markdown (MD) format (see Section 3.2). Since every dataset has its own metadata, it ensures a precise and long-lasting documentation. Another benefit is that pictures, which document the spectral conditions, can be integrated and then clearly assigned to each dataset.

The necessary groups of information for each metadata are personal information, project description, field-log & spectral documentation, and an explanation of each column from the corresponding dataset (see Supp. Figure S1, Table S2).

### 3.2 Data formats and file naming

The SPECTRE data framework is, at its core, based on two common data formats namely Comma-separated Values (CSV) for the observation tables and Markdown (MD) for the metadata. These common data formats are supported by different operating systems and guarantee versatility as well as a long-lasting data usage. Besides these core data formats (CSV and MD) other formats like Tagged Image File Format (TIFF) for geospatial maps or Data File (DAT) are possible. Within the third branch of the data framework (see Figure 3, ‘*III_db_final*’) only CSV and MD are found, as these two formats are meant to be the outcome of data processing pipelines.

The naming of the files itself is defined by a simple combination of acronyms as well as a corresponding date. For example, ‘UNS_PIK_20220504’ where UNS represents an abbreviation for the sensor used, PIK is the abbreviation for the study site, and the last number represents the date of the measurement. When multiple acquisitions are performed using the same sensor at the same site on the same date, the acquisition time (HHMM) or an additional acquisition identifier can be appended to the filename to ensure uniqueness and facilitate the identification of specific measurement sessions. Besides the simplicity of the file naming, another benefit of naming the files as proposed is that it includes the hierarchical folder structure. This ensures in a case of any loss of the folder structure that the files can be easily reconstructed into the hierarchical structure.

### 3.3 Workflow

One important principle of the SPECTRE data framework is its flexibility (see Chapter 2). The data framework, with its simple design (see Section 3.1) and the use of straightforward data formats (see Section 3.2), makes it easy to expand collections e.g. by incorporating new projects, new sensors, new datasets. Furthermore, its structure and reliance on text-based files also make it easy to implement automated checks that regularly verify the integrity of the data framework. Therefore, the workflow presented here starts with the implementation of the experiment into the experimental design and ends with data storage (see Figure 5). While the single steps shown in Figure 5 should be followed, the presentation of the processing can be also done in a different way and what is shown here is more of an example.

**Figure 5:**
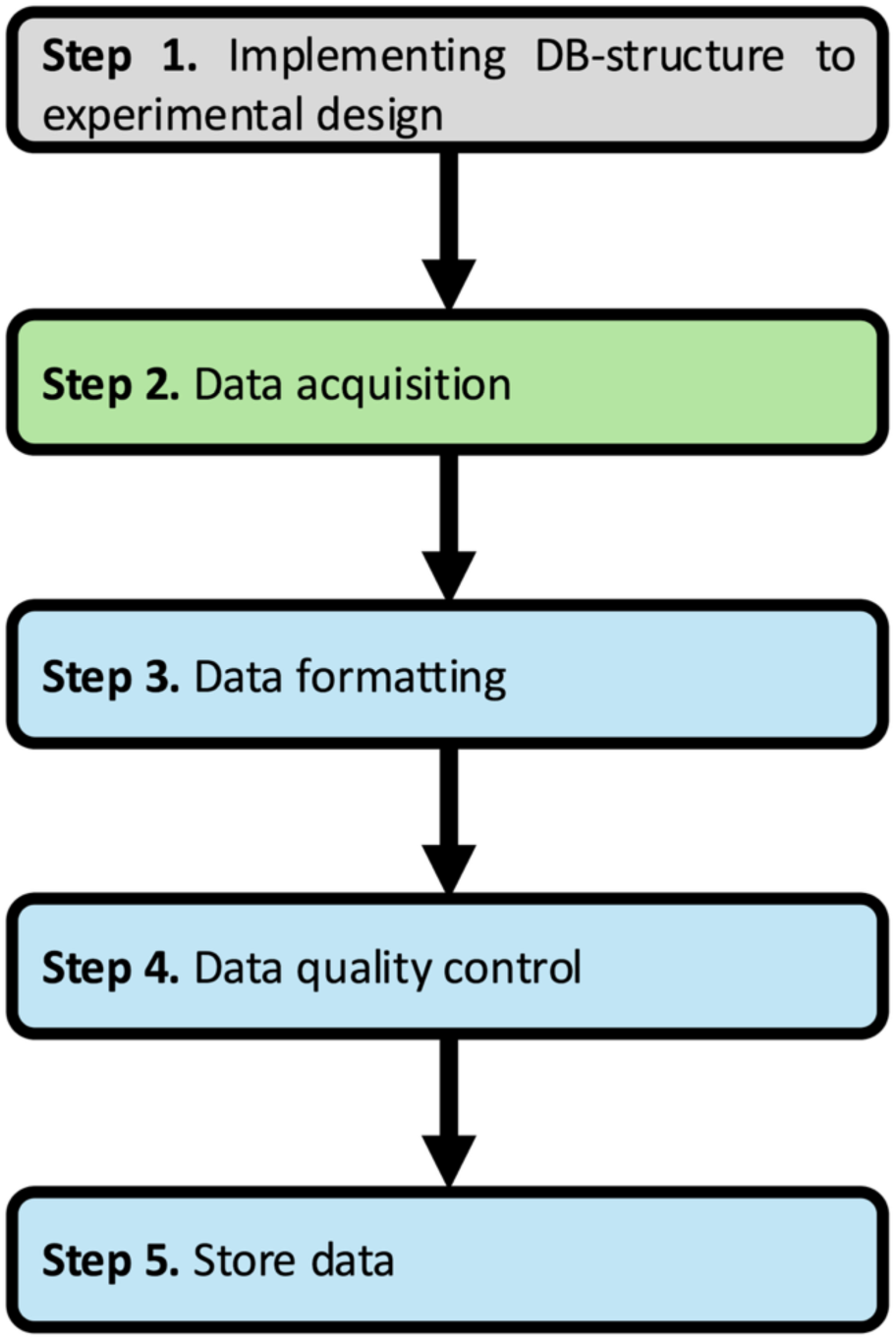
Workflow with five steps from planning (Step 1. grey), to data acquisition (Step 2 green) to data processing and storing (Step 3-5 blue).

The implementation of the data framework design into the experiment ahead is an important step, as it assures and simplifies the later data analysis. This starts by using the primary key (PK) and addressing it to a single tree (see Chapter 3.1.2). Also, the implementation of LTK at this point will ensure lower redundancy while keeping all necessary information. Lastly the metadata can be predesigned by writing out the project objectives and explaining the final columns of each variable. When these preparations have been done, the actual data acquisition can start.

After data acquisition, the actual work for adding data to this data framework starts as the data from different sensors need to be processed (see Figure 5, Step 3-5). Before starting the processing of data, the data should be stored within the *‘I_db_raw’* and the work should be done with a copy of these raw data. The goal of dataset processing is the transformation the data format into a CSV file and to establish and create the correct columns and dataset layout. Also, in case of any calculations, e.g. for vegetation indices or atmospheric vapor pressure deficit (VPD), this should be part of the processing step. All the steps can be done with scripts coded in e.g. R or Python. For easy handling the first step should be a transformation of the datasets into a CSV file, ensuring that the observations are rows and the variables are columns. When such a transformation has been done, the addition of a primary key is the next step. Therefore, mostly a lookup table is used which corresponds to a shorter, fieldwork-friendly usable identifier (ID). At next the time column should be also formatted to the format of *‘YYYY-MM-DD HH:MM:SS’*. When done, calculations can be made and new columns are added. Also, another step here is the initial quality control of the data observations, where bad observations should be removed (these occur when a measurement has been taken wrongly, is stored on the device and therefore must be removed afterwards. These ‘bad observations’ must be documented). Lastly, it is important to note that most of these steps can be combined within one loop of code, multiple datasets from a single sensor can be processed by a single block of code so ultimately or ideally for each sensor a pipeline must be coded once and can be used for repeatedly.

Following the data processing the data quality control is the next important step. The goal here is to ensure a standard quality and this step provides a chance to link outliers with certain problems that occurred while sampling (e.g. if spectral data in March look significantly different to other data and the metadata note that there has been snow it might be worthwhile to mention this). In general, it is not recommended to remove outliers mathematically, as this should be done by the end-user. For the data framework the data should stay in a rather plain form because one isn’t sure whether a mathematical outlier might be biologically relevant. What is an important quality control step is to visualize the data in a very simple way. Visual data control provides an intuitive impression of data quality that facilitates interdisciplinary discussion and strategic planning, while dynamic visualization of included values, variables, records, and absent combinations yield an invaluable summary to ensure comprehensive accounting of missing data (R. Ruddle, 2023, 2026).

## 4. Data quality

Due to its structure, the SPECTRE framework is designed to allow the automation of a set of procedures for data quality evaluation, standardization, and monitoring of dataset integrity. These verification procedures are intended to be performed when new data are added to the dataset, during their preprocessing, and during their final transfer to the III_db_final folder. The objective of this methodology is to ensure that the data integrated into SPECTRE follow a common structure and common conventions (see Chapter 3), in order to facilitate their reuse, analysis, and integration into automated pipelines, notably for artificial intelligence methods.

In parallel, standardized quality criteria can be applied to the different types of data, allowing users to more easily assess their reliability and compare analyses using a common set of criteria.

The verification procedures can therefore be separated into several categories. Standardization procedures make it possible to verify that the new data respect the conventions defined by SPECTRE. These verifications are particularly important when new data are added to the dataset. Due to the structure of SPECTRE, they can be automated from the submission of the data, following an approach similar to continuous integration (CI) mechanisms used with version control systems (such as Git), where a series of tests is executed before new data are integrated into the main structure. These tests can, for example, verify file naming, the presence of expected columns, date formats, associated metadata, as well as the consistency of measurement units and variable definitions (e.g., water potential expressed in bar or MPa). This element is even more useful because, during data collection campaigns, multiple researchers or data collectors can be involved and can generate a variety of data over relatively short periods of time. This context increases the risk of data-entry errors, inconsistencies between users, as well as misunderstanding or incorrect application of the structuring conventions specific to certain sensors or types of data.

Integrity checks, in contrast, aim to verify that the addition or modification of data does not compromise the overall consistency of the data framework. They can notably verify the validity and uniqueness of identifiers, the correspondence of primary keys with lookup tables, the absence of duplicates, or the consistency of the relationships between the different datasets. These verification procedures are intended to be performed regularly when data present in the dataset are modified, during preprocessing procedures.

Finally, quality checks within the SPECTRE data framework aim to characterize the quality of the measurements themselves according to criteria adapted to each type of data. One of the particularities of a framework such as SPECTRE is that it is designed to contain data collected across different scientific disciplines, such as hyperspectral images, soil moisture measurements, or pigment concentrations. The use of these data can therefore be complex because of their naturally multidisciplinary nature and because their quality can be evaluated according to very different criteria.

Standardizing quality metrics and systematically incorporating them during the integration of data into the data framework are therefore important. They make it possible to provide users, independently of their field of expertise, common indicators to understand data quality. Quality metrics are not necessarily generalizable across different sensors, as data quality cannot always be assessed using the same criteria. However, they should be stored alongside the corresponding datasets in CSV format, accompanied by documentation describing their meaning, calculation methods, and expected value ranges.

An important element of the SPECTRE structure is that no data stored in I_db_raw, except in exceptional cases, are deleted when a problem is detected. Affected data are instead flagged in order to warn users about their quality or their possible limitations, while keeping a trace of the original data and of the decisions made during their processing. Indeed, some noisy or atypical observations, including statistical outliers can nevertheless contain useful or biologically relevant information. Given that artificial intelligence algorithms can be sensitive to biases and outliers in training data, documenting these observations is essential to help users make informed decisions about their inclusion or exclusion when investigating biological processes.

## 5. Trackability

The organization of the SPECTRE framework into three folders was designed to allow the implementation of data traceability routines. These three folders correspond to the main phases of the data life cycle: its collection, its processing, and its availability to users (see Figure 5). The standardization, for each dataset, of Markdown files containing information on the origin and structure of the data (source filename, software version, processing script etc.), coupled with the automation of verification procedures, makes it possible, when necessary, to trace the path of a final data item back to its original raw data.

The structure also facilitates the integration of versioning and modification logging mechanisms in order to keep a record of the different transformations applied to the data throughout their life cycle. These mechanisms can record the date of a modification, its nature, the script or procedure used and, when relevant, the user responsible for the modification. However, these elements remain optional depending on the needs of the data to be stored, since they add complexity to the collection process that can sometimes be incompatible with the data collection workflow or with the technical capabilities of the teams adopting a SPECTRE storage structure.

## 6. Interface

The current prototype (insert URL here) uses a lightweight client-server architecture consisting of a backend built with FastAPI (Python) (https://fastapi.tiangolo.com/) and a JavaScript frontend built with the Tabulator data grid library (https://www.tabulator.info/). The frontend dynamically loads and displays CSV datasets served through FastAPI endpoints, enabling users to browse, filter, sort, and search large collections of metadata directly in the browser. The CSV files used for web-based exploration have been condensed from the original datasets (generated within the FastPheno project) to include only the metrics referenced in the associated markdown documentation. As a result, individual files are typically between 1 MB and 5 MB in size, providing fast load times and responsive filtering within Tabulator. To generate Tabulator-compatible datasets, CSV files should contain dedicated columns for all desired filterable attributes, including identifiers such as the full file path to flight data, acquisition date, analysis type, location, and other relevant metadata. Additional content, such as linked sky imagery capturing cloud cover on a given UAV flight day, can be incorporated through URL columns or explicit action fields that open associated image files (.png, .jpg, etc.) in a separate viewer. Future enhancements include a submission portal containing downloadable CSV and markdown templates, along with clear instructions for contributors, allowing standardized metadata collection while avoiding the need for direct file uploads to the server and reducing exposure to potential security risks.

## 7. Data Demonstration

To date, the SPECTRE data framework has successfully ingested and harmonized multi-source datasets spanning two distinct field monitoring sites focused on a primary target tree species white spruce (*Picea glauca*) (see Table 1). Across this operational window (2022–2024), the repository aggregates data from 25 UAV flight campaigns covering a total of 12,000 individual trees, yielding 131,984 high-resolution tree reflectance curve observations. Complementary ground-truth measurements comprise 7,799 combined leaf-level spectral reflectance and chlorophyll fluorescence observations, alongside detailed edaphic characterization including 59 soil texture records and high-frequency soil moisture profiles. Furthermore, local meteorological records provide continuous temporal context, encompassing 21,900 daily aggregated values and 52,608 hourly observations.

**Table 1:**
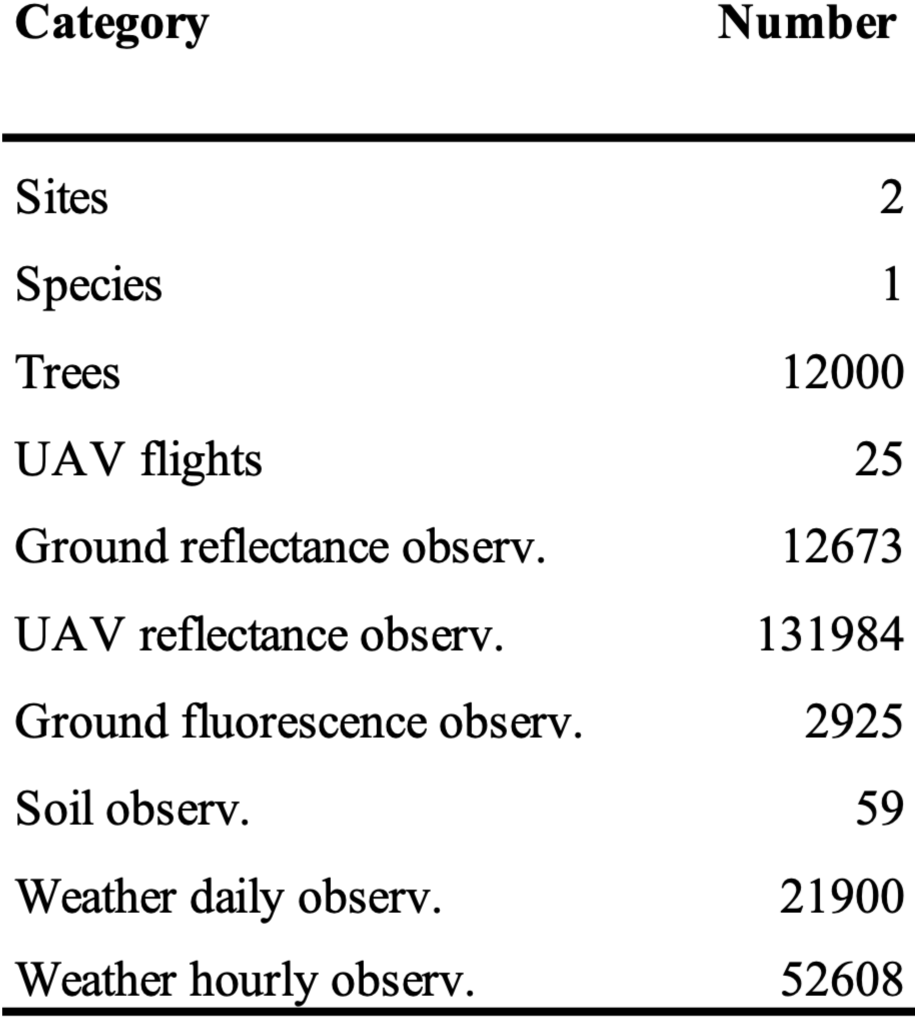
Categories of SPECTRE framework features which describe from how many Sites and how many species are measured, the total amount of trees, how many flights have been done and how many observations for a given category are available.

By implementing a robust relational database architecture structured around a standardized primary key, all disparate data streams are seamlessly linked. As illustrated in Figure 6, this framework facilitates direct correlation analyses between ground-based measurements and UAV-retrieved spectral indices, which can be executed dynamically at the individual tree, site-specific, or macro-aggregate scale. Furthermore, a distinctive capability of the architecture is the direct temporal synchronization of high-resolution meteorological time series with airborne remote sensing datasets (Figure 7). This integration enables immediate evaluation of environmental drivers on canopy-level physiological status without intermediate formatting bottlenecks.

**Figure 6:**
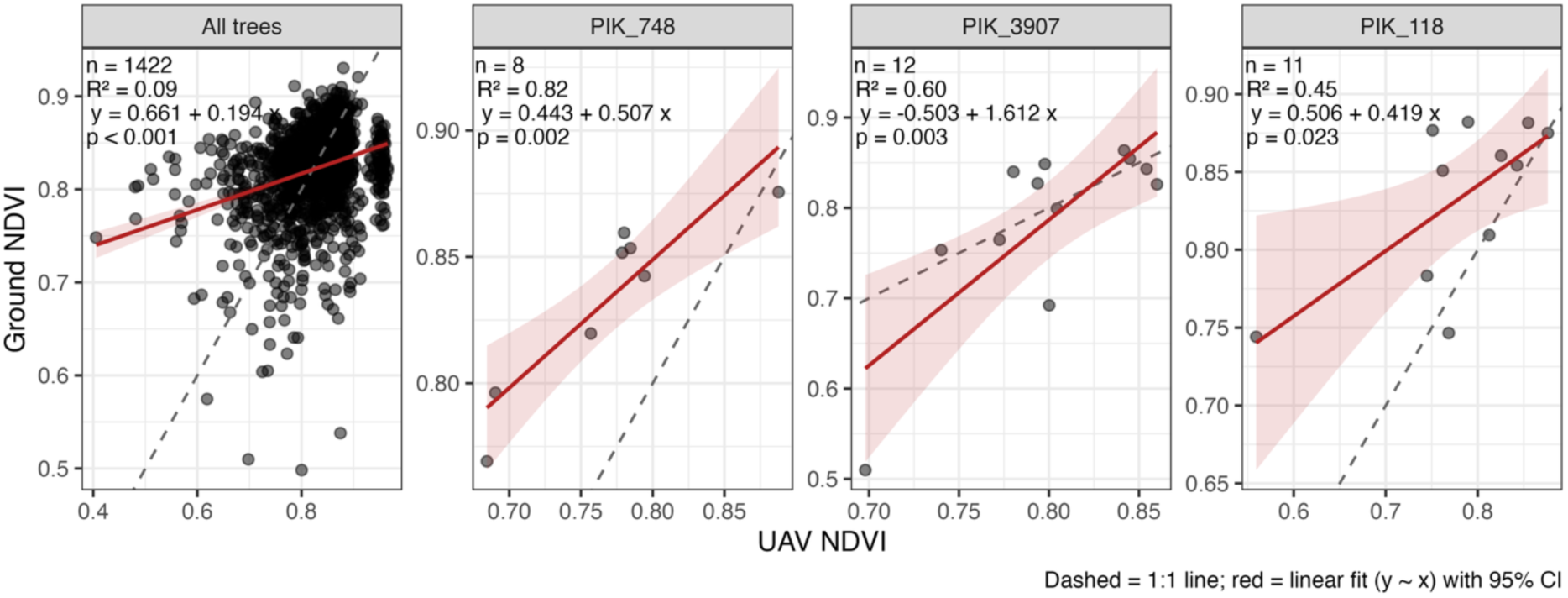
Linear regression analysis comparing ground-measured Normalized Difference Vegetation Index (NDVI) against UAV-retrieved NDVI. Results are displayed for the aggregated population (“All trees”, *n* = 1422) as well as representative individual trees (PIK_748, PIK_3907, and PIK_118). Solid red lines represent the ordinary least squares linear fit (*y* ∼ *x*) with corresponding 95% confidence intervals (shaded pink regions), while dashed gray lines indicate the 1:1 reference relationship. Statistical parameters including sample size (*n*), coefficient of determination (*R*^2^), regression equations, and *p*-values are embedded within each panel.

**Figure 7:**
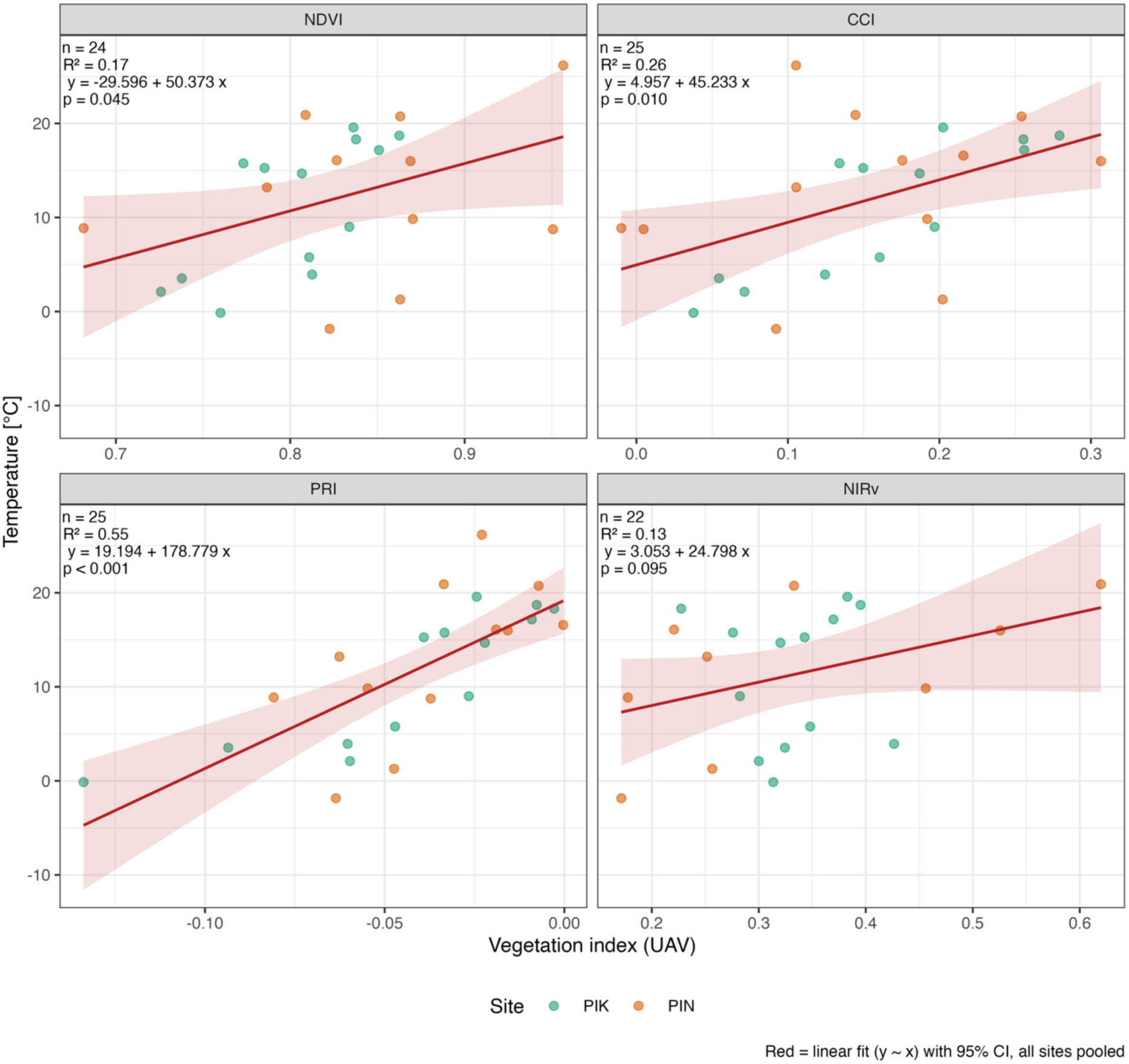
Linear regression relationships between meteorological air temperature (°C, daily value) and UAV-retrieved vegetation indices Normalized Difference Vegetation Index (NDVI), Chlorophyll/Carotenoid Index (CCI), Photochemical Reflectance Index (PRI), and Near-Infrared Reflectance of vegetation (NIR*_v_*) pooled across sites (PIK in green and PIN in orange). Solid red lines represent the ordinary least squares linear fit with 95% confidence intervals (shaded pink regions). Statistical annotations per panel indicate sample size (*n*), coefficient of determination (*R*^2^), regression equations, and *p*-values.

In operational terms, this unified data management workflow fundamentally streamlines traditional ecophysiological data processing. Tasks that historically required hours of manual data wrangling across fragmented formats and proprietary structures can be replaced by simply uploading the framework into any program e.g. RStudio, Python. By eliminating administrative friction and establishing a direct pathway from raw sensor streams to analysis-ready matrices, the SPECTRE framework demonstrates that high-throughput, multi-scale data integration can be achieved with minimal user intervention, enabling researchers to focus intensively on interpretation and ecological modeling.

## 8. Conclusion

The SPECTRE data framework is, to our knowledge, the first framework that systematically integrates data storage across the fields of plant ecophysiology, remote sensing, and micro-environmental values. By bridging the long-standing gap between fine-scale ground measurements (e.g., photosynthetic parameters and leaf water potential) and high-resolution aerial imagery, SPECTRE enables quick analysis while ensuring easy data discoverability and usability through an interface and basic data formats. It is therefore particularly well suited to small research teams due to its lightweight, easy handling, and flexible architecture.

To achieve long-term sustainability without continuous server or software expenses, SPECTRE intentionally avoids complex relational database management systems in favor of a simple file- and-folder structure built on open formats like Comma-Separated Values (CSV) and Markdown (MD). Supported by a clean, intuitive interface, this simplicity facilitates that datasets remain easily accessible and searchable across research teams and over time. Consequently, SPECTRE provides a robust, low-cost defense against “data entropy”, the gradual loss of data accessibility driven by the established long-term infrastructure (Michener et al., 1997; Pepe et al., 2014; Vines et al., 2014).

Beyond data preservation, SPECTRE directly adds value to the critical training data bottleneck that currently limits the application of artificial intelligence (AI) and deep learning in forestry. By establishing a unified primary key, the framework links individual-tree crown segmentation (ITCS) masks directly with co-registered spectral signatures and physiological ground truths (Colomer et al., 2026). This standardized structure improves data consistency and integration, streamlines automated workflows, and prepares raw multi-sensor datasets for immediate ingestion into advanced machine learning pipelines.

Ultimately, SPECTRE offers an adaptable, low-cost data management protocol tailored for individual remote sensing and ecophysiology laboratories, as well as broader collaborative research networks. As climate change continues to increase the vulnerability of global forest ecosystems, bridging multi-scale observations through standardized, open-access protocols becomes essential. By providing an accessible, scalable, and future-proof foundation for data synthesis, SPECTRE empowers the ecological community to advance proactive, data-driven strategies for climate-smart forestry and sustainable ecosystem management.

## Acknowledgements

The authors gratefully acknowledge the financial support granted by Ontario Genomics, Genome Quebec, and Genome Canada. We are thankful to the people involved in the FastPheno project acquiring the data on which this framework has been developed on. Furthermore, we are thankful for the engaging discussions at the FastPheno annual meetings as well as the regular meetings on data organization. Special thanks go to Vincent Segnier, who was responsible for the UAV flights at Pintendre and provided particularly valuable feedback on data organization. Felix Tran and Vincent Lau also deserve special recognition for their work on the interface, as well as their input on data organization.

## Author contributions

VPR, SC, and IE designed the data framework. VPR, JG, and SC processed the data. VPR created all the figures and wrote the first manuscript, with parts written by SC and NP. All authors intensively discussed and revised the manuscript.

