## Supplementaries for "SPECTRE: A versatile data framework for integrating spectral, environmental, and ecophysiological trait data"

### Supplements

#### *Tables supplements*

**Table S1**: Abbreviations

| Abbreviation | Explanation |
|---|---|
| PK | Primary key |
| CSV | Comma Separated Value |
| MD | Markdown |
| SPECTRE | Spectral, Trait, and Remote-sensing Ecophysiology database |
| LKT | Lookup tables |
| TIFF | Tagged Image File Format |
| DAT | Data File |
| ID | Identifier |
| AI | Artificial Intelligence |
| DL | Deep learning |
| ML | Machine learning |
| SQL | Structured Query Language |
| HDF5 | Hierarchical Data Format, Version 5 |
| NetCDF | Network Common Data Form |

**Table S2**: Examples of metadata fields used to document hyperspectral and LiDAR data collected as part of the FastPheno dataset, following the structure of the SPECTRE framework. The fields are organized into five categories covering general mission information, environmental conditions, UAV and sensor configurations, flight parameters, and data traceability. The metadata are stored in YAML files to facilitate access, readability, and automated processing.

| **Category** | **Metadata fields** |
|---|---|
| ***General information*** | Project name, mission identifier, acquisition date, study site, supervisor, operators, observers |
| ***Environmental conditions*** | Weather conditions, wind speed, wind gusts, temperature, illumination conditions |
| ***UAV and sensor configuration*** | UAV platform, sensors, sensor mounting, positioning system |
| ***Flight parameters*** | Flight number, altitude, speed, ground resolution, frame period, exposure time, flight management software, takeoff and landing times |
| ***Data traceability*** | Recorded events, battery identification, GNSS corrections, control points, raw data storage location, additional comments |

***Figures supplements***

### UniSpec Reflectance Metadata

#### Personal Information

| Function | Name | Email | Phone |
|---|---|---|---|

#### Project Information and Operation Comments

| Item | Details |
|---|---|
| Description: | |
| Operation comments: | |

#### Metadata Description

| column_name | description |
|---|---|
| primary_key | Unique identifier consiting of site (e.g. PIK) and the trees rolling number (e.g. _1001). |
| datetime_est | Date and time of measurement acquisition (EST timezone). |
| datetime_ust | Date and time of measurement acquisition (UST timezone). |
| date | Date of spectral acquisition (YYYY–MM–DD). |
| time | Time of spectral acquisition (HH:MM:SS). |
| campaign | Field campaign identifier or campaign number. |
| sample | Sample number within the tree or measurement sequence. |
| acquisition_time | Acquisition duration or integration time of the measurement in seconds. |
| tree_key | Combined unique key identifying the tree and acquisition event. |
| orientation | Cardinal orientation of the measurement relative to the tree canopy. |
| measurement_number | Measurement number from the device. |
| x300 to x1000 | Reflectance wavelength from 300 to 1000 nanometer. Interpolated to 1 nm wavlength. |
| PRI | Photochemical Reflectance Index; Formular: (‘531‘ – ‘570‘) / (‘531‘ + ‘570‘) |
| NDVI | Normalized Difference Vegetation Index; Formular: (‘800‘ – ‘630‘) / (‘800‘ + ‘630‘) |
| NIRv | Near–Infrared Reflectance of Vegetation; Formular: NDVI * ‘860‘ |
| CCI | Chlorophyll/Carotenoid Index; Formular: (‘532‘–‘630‘) / (‘532‘+‘630‘)‘ |
| ChapA | Chlorophyll fluorescence ratio; Formular: 675‘/‘700‘;Estimated chlorophyll a content. |
| ChapB | Chlorophyll fluorescence ratio; Formular: 675‘/(‘700‘ . ‘650‘); Estimated chlorophyll b content. |
| PSRI | Plant Senescence Reflectance Index;Formular: (‘678‘ – ‘500‘) / ‘750‘ |
| SIPI | Structure Insensitive Pigment Index; Formular: (‘800‘ – ‘445‘) / (‘800‘ – ‘680‘) |
| NDRE | Normalized Difference Red Edge Index; Formular: (‘790‘–‘720‘) / (‘790‘+‘720‘) |
| WBI | Water Band Index; Formular: ‘900‘ / ‘970‘ |
| CIRE | Chlorophyll Index Red Edge; Formular:(‘750‘ / ‘710‘) – 1 |
| EVI | Enhanced Vegetation Index; Formular: 2.5 * (‘858‘ – ‘645‘) / (‘858‘ + 6.0 * ‘645‘ – 7.5 * ‘469‘ + 1.0) |
| EVI2 | Two–band Enhanced Vegetation Index; Formular: 2.5 * (‘858‘ – ‘645‘) / (‘858‘ + 2.4 * ‘645‘ + 1.0) |
| SAVI | Soil Adjusted Vegetation Index; Formular: ((‘860‘ – ‘660‘) / (‘860‘ + ‘660‘ + 0.5)) * (1 + 0.5) |
| GNDVI | Green Normalized Difference Vegetation Index; Formular: (‘800‘ – ‘550‘) / (‘800‘ + ‘550‘) |

**Figure S1:** Example metadata organized by ‘Personal information’, ‘Project Information and Operation Comments’, and ‘Metadata Description’. The latter explains every column of the corresponding CSV dataset with the actual observations.
